# Impairment of zebrafish reproduction upon exposure to TDCPP

**DOI:** 10.1101/176909

**Authors:** Xiecheng Liu, Guofeng Jia

**Affiliations:** Department of Health Management, Qingdao Municipal Hospital, Qingdao 266021, Shandong, China

**Keywords:** TDCPP, Reproduction, Zebrafish

## Abstract

TDCPP is one of the most common organophosphate flame retardant, which has been widely used in many products. It has been detected in the environment and biota; however, its potential affect to the wildlife and human health remains largely unknown. In this study, we aimed to investigate the effect of long-term exposure to TDCPP on fish reproduction. Zebrafish eggs were treated with various concentration of TDCPP (0, 1, 10 and 100 μg/L) from 1 day post-fertilization (hpf) to 6 months. The fecundity of female fish was significantly decreased as indicated by reduced embryos production. The egg quality was decreased and the malformation rates were increased in the F1 generation. Taken together, long-term exposure to TDCPP affects the reproduction of zebrafish.

## Background

Tris(1,3-dichloro-2-propyl) phosphate (TDCPP) is an orgnophosphate flame retardant, which has been widely applied in the products of resins, latexes and polyurethane foams in the furniture [1-5]. TDCPP is easily to be released into the environment because its non-chemically bounding property to the household items [3, 6-10]. Thus, TDCPP can cause pervasive contamination in the air, surface and ground water [11-15]. TDCPP is relatively stable in water because it is lipophilic and therefore hard to be decomposed in the environment and could lead to bioaccumulate in the aquatic animals [4, 16-18].

Previous studies have reported that TDCPP could affect the sex hormone levels in human cells and induce the transcriptional expression of estrogenic receptors and related genes in zebrafish [19]. TDCPP affects the estrogenic activity and interrupts endocrine levels in fish [20-22], however, the effect of TDCPP on the reproductively in TDCPP exposure on zebrafish reproduction, especially in a long-term exposure period, which most reflects the affection of TDCPP pollution in the aquatic environment.

## Materials and methods

### Chemicals

TDCPP, [(1,3-dichloro-2-propyl) phosphate] (analytical standard) was purchased from Sigma Aldrich. Stock solutions were prepared in dimethyl sulfoxide (DMSO, purity > 99%) and stored at − 20 °C.

### Zebrafish maintenance and TDCPP treatment

Wild type zebrafish were raised as described previously [23-27]. The fish were kept at 28 ± 0.5 °C in a 14 h light/10 h dark cycle. Fish were naturally crossed and eggs were collected. Five hundred fertilized zebrafish embryos were treated with TDCPP (0, 1, 10 and 100 μg/L) in 1-L tanks. The zebrafish larvae were put into 2.8-L aquariums at 20 dpf. Each exposure groups and controls had three replicates and received 0.001% (v/v) DMSO. Fish were mated in groups weekly to evaluate the reproductive capacity. Fecundity was calculated as the cumulative average number of embryos produced per female each time.

### Measurement of hormone concentrations

The estradiol and testosterone levels were examined in the blood samples of TDCPP-treated zebrafish as previously described [28-30]. In brief, fish were euthanized in MS-222 (200 mg/L), the blood samples from caudal vein were collected, centrifuged at 5000 × g for 10 min at 4 °C. The supernatant was collected for hormones extraction.

### Data analysis

All analyses were carried out using SPSS 18.00 (SPSS, Chicago, IL, USA). Differences between the control and TDCPP-treated group were assessed by one-way analysis of ANOVA followed by a Tukey’s test. All data were presented as the mean ± SE, P< 0.05 was considered statistically significant.

## Results

### Toxicological effects of TDCPP exposure to zebrafish

The hatching, malformation and survival rates of the F1 generation were recorded from 1 dpf to 5 dpf. There was no significantly difference in the F1 eggs derived from the P0 fish exposed to 1 and 10 μg/L TDCPP. However, the malformation rate in the F1 eggs derived from P0 fish that were treated with 100 μg/L TDCPP was significantly increased (Data not shown).

The survival rates in the TDCPP-exposed P0 fish had no significant difference during the six months of treatment. As shown in Fig. 1, the average numbers of embryos produced were significantly decreased in the groups treated with 10 and 100 μg/L TDCPP compared to the controls.

**Figure 1.**
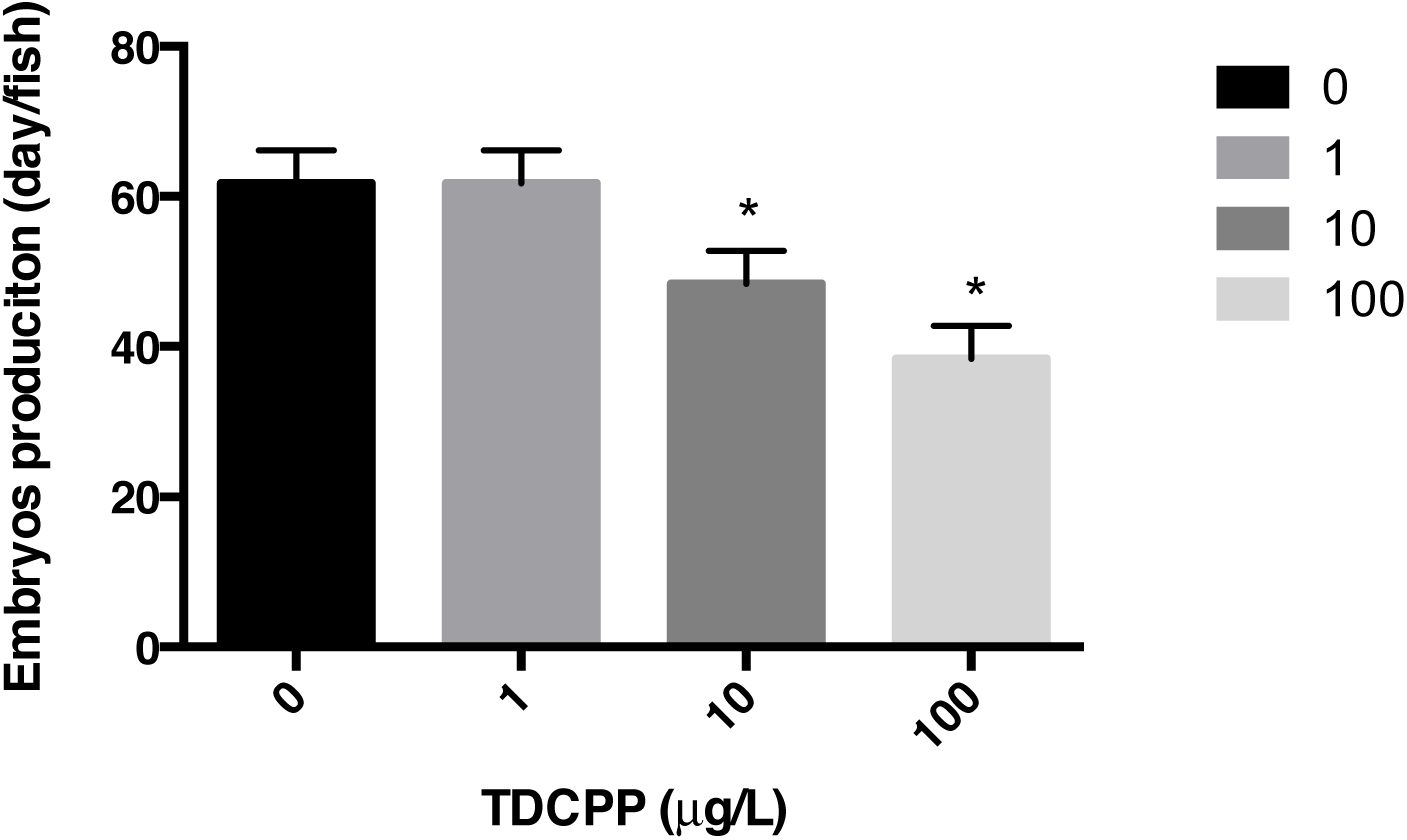
The productions of embryos in zebrafish treated with TDCPP in different concentration. *p< 0.05 present significance between exposed group and the controls.

### Hormones levels in TDCPP exposed fish

Compared to the controls, the estradiol (E2) and testosterone (T) levels in female fish treated with 10 and 100 μg/L TDCPP were increased (Fig. 2); however, the E2 and T levels in the male fish treated with TDCPP had no significant changes (Fig. 3).

**Figure 2.**
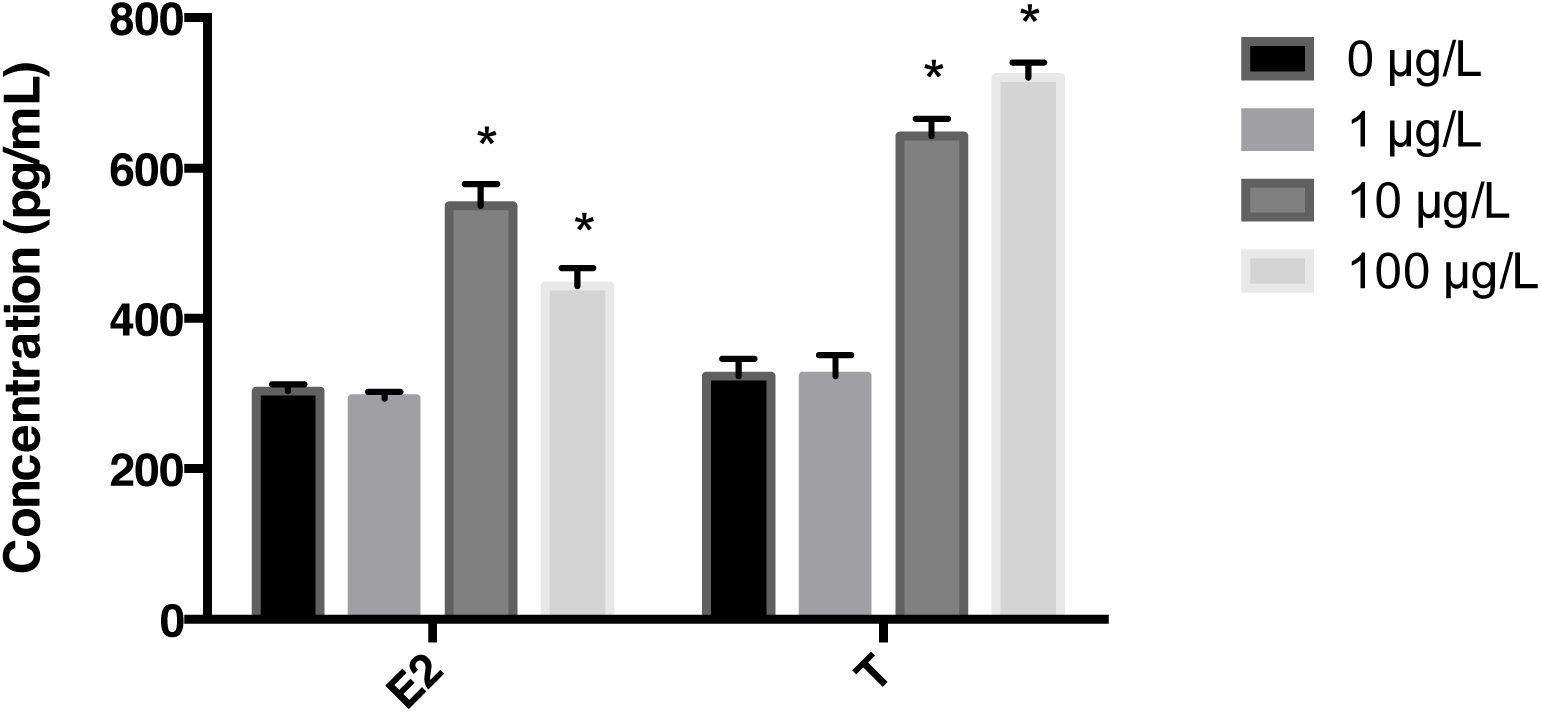
The estradiol (E2) and testosterone (T) contents in the female fish treated with TDCPP. *p< 0.05 present significance between exposed group and the controls.

**Figure 3.**
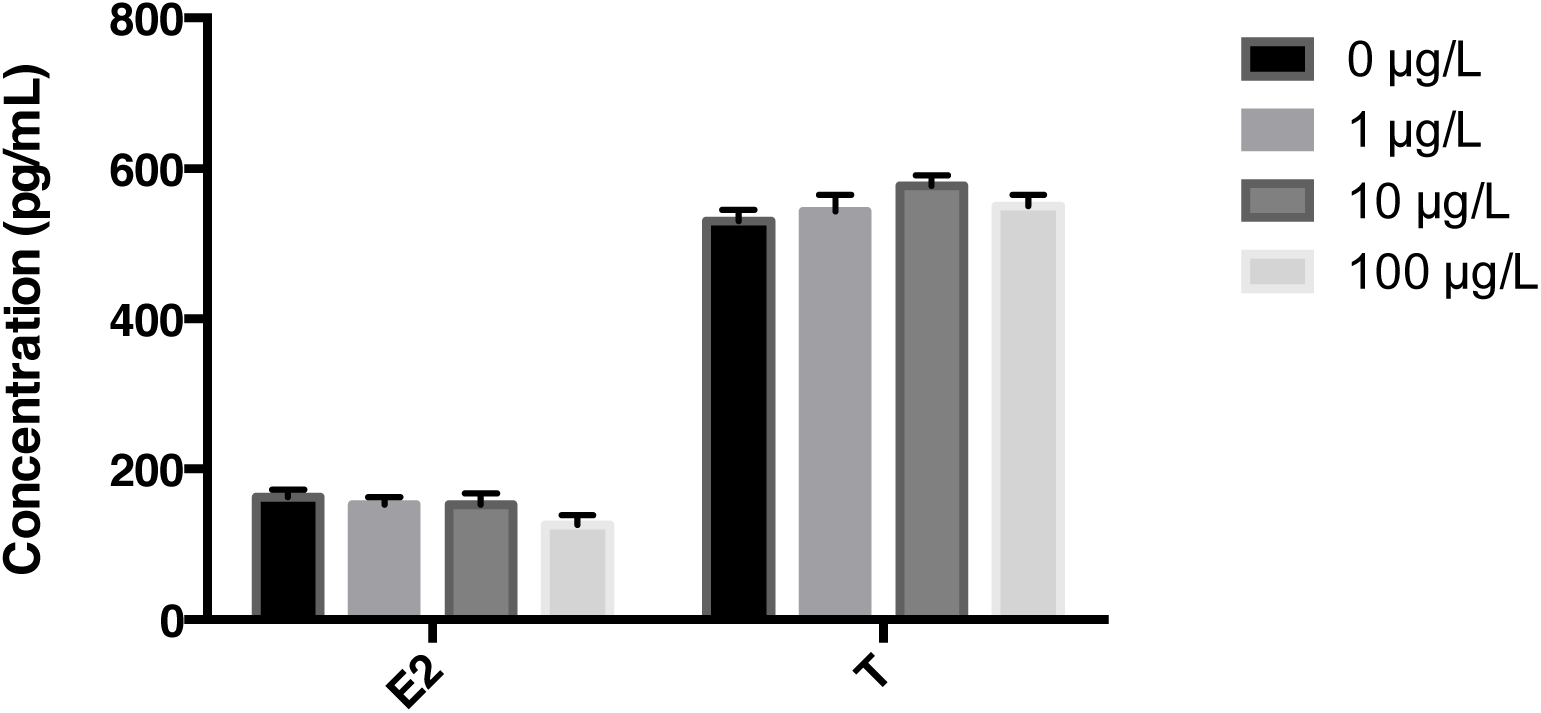
The estradiol (E2) and testosterone (T) contents in the male fish treated with TDCPP.

## Discussion

In this study, we investigated the effects of long-term exposure to TDCPP on plasma hormone levels in zebrafish, which is consistent with the previous studies that report TDCPP alters steroidogenesis in fish [31-35]. In addition, we observed that exposed to high concentration of TDCPP reduced the embryos production and caused higher malformation rates in the F1 eggs. There results confirmed that long-term exposure to environmental concentrations of TDCPP affect the fish reproductive system [35-40].

We also found that the E2 and T levels were higher in the female but not in the male fish exposed to TDCPP, which indicated that TDCPP might have gender-specific effects on the zebrafish steroidgenesis. These data were in consistent with the previous studies that endocrine disrupting chemicals have estrogenic activity and could affect the sex hormones in fish in a gender-specific manner [41-48].

The toxicity of TDCPP treatment was investigated in P0 and F1 zebrafish. We observed the malformation rate in F1 embryos that derived from P0 fish treated with TDCPP was increased. This data suggested that TDCPP had developmental toxicity to the offspring when the parental fish were treated with TDCPP. Thus, our results suggested that long-term exposure to environmental concentration of TDCPP to P0 fish resulted in developmental toxicity to the F1 embryos.

Taken together, our study confirmed that chronic treatment with environmental concentrations of TDCPP impaired zebrafish reproductive system [5, 32, 35, 38, 49, 50]. However, the effects of TDCPP on the sex hormone levels in fish still need further investigation.

